# Twisting to freedom: The evolution of copulation termination techniques across 48 species of sepsids (Diptera, Sepsidae)

**DOI:** 10.1101/2021.06.30.450518

**Authors:** Mindy Jia Min Tuan, Diego Pitta Araujo, Nalini Puniamoorthy, Jeremy M Woodford, Rudolf Meier

**Affiliations:** National University of Singapore, Department of Biological Sciences, SINGAPORE

**Keywords:** Copulation separation, disengagement, mating behaviour, evolution, copulation

## Abstract

Studies of insect mating behaviour usually focus on what happens before and during copulation. Few pay close attention to the actions needed to end copulation. However, genital separation after copulation is likely to be an important cause of mechanical stress and injuries because it often involves the withdrawal of heavily armoured male intromittent organs from membranous female reproductive tracts. Difficult and/or slow separations can also reduce male and female fitness by increasing their exposure to predation. We here report the results of a comparative study of separation behaviour in 48 species of Sepsidae (Diptera) and one outgroup. We find a surprising amount of qualitative and quantitative behavioural variability within and between species. We characterize and reconstruct three types of behaviours: 1) The sepsid ancestor likely used ‘back-off; a gentle separation technique that does not involve any pulling or twisting (https://youtu.be/EbkJvOaubZ0). 2) This potentially gave rise to the most common ‘pull’ technique where the male turns 180 degrees and pulls in an opposite direction from the female (https://youtu.be/oLf4xGpkk1s). This separation can be quick and straightforward, but in some species the ‘pull’ is slow and protracted and we routinely find dead males and/or females attached to their living partners in the latter (difficult: https://youtu.be/MbYPbXN6jr0; failure: https://youtu.be/leTiXefFzCc). 3) Finally, several species use ‘twist’, a new technique where the male rotates >360 degrees from the initial mounting position (https://youtu.be/WMUXbIPyLbk). We document that species capable of using ‘twist’ have shorter and less variable separation times than those using “pull”. However, many species capable of ‘twist’ also retain the ability to use ‘pull’ (‘back-off’/’pull’= 8%; ‘pull’ only= 41%; ‘twist’/ ‘pull’= 24%; ‘twist’ only = 27%). Overall, our study suggests that separation behaviour can vary among closely related species and highlights the significance of studying variable behavioural traits in a phylogenetic context.

## Introduction

Males and females of many insect species perform elaborate courtship behaviours before and during copulation, and many of these behaviours are species-specific (Mayr 1963; Spieth 1974; Boake 1989; West-Eberhard 1984; Eberhard 2001; Puniamoorthy et al. 2008; 2009). However, there is comparatively little information on how copulations are terminated. Most publications barely mention separation behaviour [e.g. “spontaneous disengagement”, “the male separated from the female” or “the male withdrew his aedeagus” (Wojcik 1969; Cook 1990; Wang et al. 1996; Bonduriansky 2003; Kamimura 2008)] and comparative information across multiple species is largely missing. This lack of attention is somewhat surprising because many intromittent organs are morphologically complex, evolve quickly (Eberhard 1985; Arnqvist and Danielsson 1999), and include features such as spines and armoured protrusions that suggest that withdrawal from the female could be mechanically difficult.

Difficult genital separations can impact individual fitness. For instance, a prolonged struggle to separate after copula is likely to increase the risk and exposure to predation (Wing 1988; Rowe 1994; Hosken et al. 1994; Kotiaho et al. 1998; Kemp 2012; Siemers et al. 2012; but see McCauley and Lawson 1986). Struggles to separate could also damage male and/or female reproductive organs (Kummer 1960; Ross 1983; Merritt 1989; von Helverson and von Helverson 1991; Crudgington and Siva-Jothy 2000; Blanckenhorn et al. 2002; Allard et al. 2006; Baer and Boomsma 2006; Kamimura 2007; Kamimura 2008). Indeed, morphological damage sustained after copulation may not even be caused by copulation but by separation instead (Ross 1983; Blanckenhorn et al. 2002) because only few studies can conclusively attribute reproductive tissue damage to the time spent in copulation (Merritt 1989; Crudgington and Siva-Jothy 2000; Allard 2006; Kamimura 2007, Okuzaki et al. 2012). For instance, some studies provide evidence of scarring in non-virgin females, and speculate that the male intromittent structures could be responsible for such damage (von Helverson and von Helverson 1991; Blanckenhorn et al. 2002; Baer and Boomsma 2006; Kamimura 2008). These “copulatory” wounds may be the result of the friction between heavily sclerotized male structures and the soft female genitalic membrane sustained largely during separation.

Dramatic cases of separation problems have been observed in a few species of sepsid flies (Diptera: Sepsidae), where the failure to separate routinely leads to male and/or female death (*Sepsis cynipsea*: Blanckenhorn et al. 2002; *Themira biloba*: pers. obs.; https://youtu.be/leTiXefFzCc). When this happens, there is a fitness cost to the surviving partner who drags the dead companion around. If the dead companions is male, the dead carcasses becomes a mating plug and prevents female remating which should allow sperm ample time to migrate to female sperm storage organs (Herberstein et al 2012). However, in sepsids, the difficulty in separation affects both sexes and usually leads to subsequent death of the surviving partner; i.e., an adaptive function is unlikely. It is this peculiar phenomenon that has prompted this systematic study of separation behaviour across 48 species of Sepsidae (Diptera). We reconstruct the evolution of separation behaviour by mapping it onto the phylogenetic tree for Sepsidae (Lei et al. 2013). We document variation in these behaviors and discuss why certain separation behaviours may be adaptive. Previous studies of mating behavior in sepsids, only summarised differences in separation behaviour as “short /quick” or “prolonged” (Puniamoorthy et al. 2009; Tan et al. 2011). Here, we elaborate on separation as with or without a 180 degree turn (Eberhard and Pereira 1996; Tan et al. 2011), and with a clock- or anti-clockwise pivot around the joined genitalia (Parker 1972). We hope our study reiterates the importance of studying detailed variation of specific behaviours across multiple species.

## Methods

We studied the separation behaviour for 48 sepsid species that were identified, reared and established in the lab following previously published methods (Ang et al 2013; Meier 2017; Puniamoorthy et al. 2009; Tan et al. 2011). Sexually mature virgin flies were paired in small Petri-dishes and their mating behaviour was recorded under a microscope for at least 35 minutes or until copulation terminated and each pair of flies was only used once. The microscope was either attached to a video cassette recorder or a digital camcorder (Canon LEGRIA HF S30). The final dataset included video recordings from previous studies (not analysed in detail for separation) as well as new information for 15 additional species (Supp. table A). In total, we conducted and recorded ca. 2000 mating trials of which 426 were successful (2 to 17 recordings per species for 49 species). For 271 trials across 33 species, we also collected information on body size of males and females (Supp. table A). Based on these video recordings, we analysed the separation behaviour frame-by-frame (25 frames/sec) and recorded the time taken in seconds and frames for a male to separate from the female, which is measured from the moment the male dismounts, to the moment the paired genitalia lose contact.

### Evolution of separation behaviour

In order to understand the evolution of different separation techniques in sepsids, a three state-character (‘back-off’, ‘pull’ and ‘twist’) was defined and optimized onto the latest phylogenetic hypothesis for Sepsidae (Lei et al. 2013) using Mesquite (Maddison and Maddison 2015). In an attempt to account for intraspecific variation, we omitted *Meroplius fukuharai* from the final analysis because only one separation video was available. One complication in analysing the evolution of separation behaviour is that many species are reluctant to mate under experimental conditions (=sample size problem) and that we observe intraspecific variability with regard to separation techniques. We therefore had to analyse the origins of ‘pull’ and ‘twist’ for different sample sizes (N=2-10) in order to identify stable results; i.e., we performed a sensitivity analysis to account for sample size and to test the strength of evidence for the evolution of ‘twist’ separation behaviour from ‘pull’.

### Polymorphic species

Many species were polymorphic with regard to separation technique. Preliminary data indicated that polymorphism was more common in species that predominantly perform a ‘twist’ instead of a ‘pull’. In order to test formally whether polymorphisms are concentrated in species that are capable of performing the ‘twist’, we coded a polymorphism character where polymorphic species were scored as ‘0’ and monomorphic species as ‘1’. We then created a separation character that ignored polymorphism and scored species according to the behaviour performed in majority of trials. Species were scored as ‘0’ if the majority of males were performing a ‘pull’, as ‘1’ if it was a ‘twist’. We scored the few species with ‘back-off’ as ‘?’. We then tested for correlation between these two variables using correlation analyses in Mesquite using the pairwise comparisons method (Read and Nee 1995; Maddison 2000), and Pagel’s method (Pagel 1994). Note that this approach involving a “polymorphism character” had to be adopted because the sample sizes were too small to code frequencies directly.

### Relationship between separation technique and sexual size dimorphism (SSD)

We hypothesized that males may be able to separate more readily through ‘pull’ if the sexual size dimorphism i.e. the difference in body sizes between males and females, was small. Conversely, the more complex ‘twist’ behaviour could be an adaptation to separation difficulties in species with larger SSD. In order to test this hypothesis, we analysed a subset of data, including species that displayed both behaviors and constructed a logistic regression model with binomial distribution and logit-link function using SSD as the explanatory variable to explain the separation technique used (‘pull’ versus ‘twist’) (McCullagh and Nelder 1997).

### Relationship between separation technique and separation time

In order to determine whether SSD and the type of separation were predictors of separation time, we used averaged values for each species in a phylogenetic generalized least squares model (PGLS: Grafen 1989; Martins and Hansen 1997), which accounts for non-independence in data points between species due to shared phylogenetic history. We scored separation type for each species according to the technique used by majority of trials. Here, we note that the presence of intraspecific variation in separation behaviour indicates that separation technique may be a phylogenetically labile trait, and there may be no need to perform a phylogenetic correction. Nonetheless, we wanted to test if there was any correlation even after accounting for shared ancestry. We began with a maximal model with all predictors and performed a backward elimination, eliminating predictors that did not significantly contribute to the model, one at a time, until we reached the final model.

### Relationship between separation technique and variation in separation time

One possible explanation for the evolution of the ‘twist’ behaviour is to increase the efficiency of separation (i.e. mean and variance). To test this, we compared the separation time of males of two groups: (1) species that were observed to only do ‘pull’ (2) species that were observed to do ‘twist’ at least once. We used the Brown-Forsythe test of variance on deviation from group medians to compare the variance between the two (Brown and Forsythe 1974). We then used a Welch test to analyse group differences in means, as the data deviated from normal distribution, and group variances and sample sizes were not equal (Welch 1951).

### Handedness

The male aedeagus in sepsids is often asymmetrical which can influence the coupling of the male and female genitalia during copulation (Araujo 2015). Males can also vary in the direction in which they perform a certain separation behaviour (clockwise versus anti-clockwise). We examined aedeagus morphology of males in each species and performed a Chi^2^ test for the relationship between technique and directionality of separation.

All tests were performed in R, ver. 3.3.1 (R Core Team 2016) using the following packages: Welch test and GLM with *stats* (R core Team 2013); Brown-Forsythe test with *car* (Fox and Weisberg 2011); PGLS with *ape* and *caper* (Paradis et al. 2004; Orme et al. 2013)

## Results

Separation behaviour in 48 species of sepsids and one outgroup was surprisingly varied and diverse. We group the behaviours into three different separation techniques and documented large variations in separation times within and across species (e.g., *Sepsis lateralis* 1.5 ± 1 s; *Nemopoda nitidula* 83.25 ± 242.80 s) (Fig. 1). We observed three types of behaviours: 1) ‘back-off’, in which males separate from females by dismounting to the side and ceasing genital contact (https://youtu.be/EbkJvOaubZ0); 2) ‘pull’, in which males dismount, turn 180 degrees away from females and pull in an opposite direction (https://youtu.be/oLf4xGpkk1s); and finally 3) ‘twist’, in which males dismount and ‘twist’ in a revolution of 360 degrees or more from the initial mounted position, using the genitalia as a hinge (https://youtu.be/WMUXbIPyLbk). ‘Twist’ occurs in both directions (clock and anti-clockwise, see section on Handedness), and ‘twist’ males differed in the number of revolutions they performed (ca. 1-3) which varied both within and across species (Fig. 1). In many species, all males performed the same behaviour, but males in 16 species exhibited intraspecific variation in separation technique. Four species performed both ‘back-off’ and ‘pull’, and 12 species exhibited both ‘pull’ and ‘twist’ but there were no species that performed all three separation techniques (Fig. 1).

**Figure 1.**
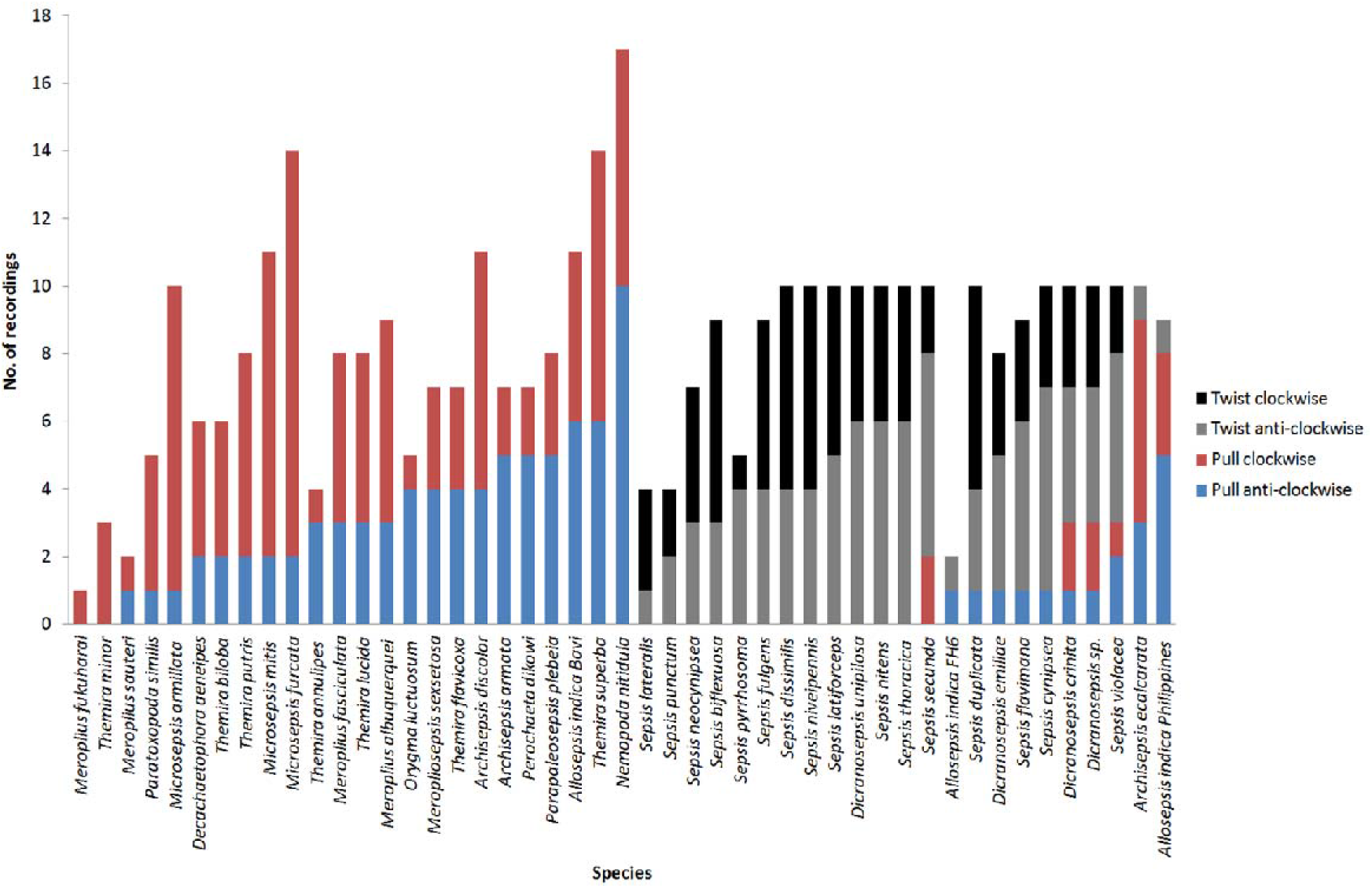
Sample size, direction of separation, and distribution of ‘pull’ and ‘twist’ across species.

### Evolution of separation behaviour

Based on our ancestral state reconstruction, there is a clear shift from ‘pull’ to ‘twist’ as a derived behaviour with multiple origins in sepsids (Fig. 2). There were three species, in which ‘twist’ was performed at a low frequency (10-33%). A particularly interesting case is *Archisepsis ecalcarata* which usually performs a ‘pull’ in a stereotypical manner, except for one male which seemed to “chance onto” ‘twist’ after his initial ‘pull’ attempt failed to break genital contact (https://youtu.be/uNPB7cURU7A). Another species with an “intermediate” behaviour is *Nemopoda nitidula* where a few males that experienced separation difficulties via ‘pull’ were also moving side to side but they did not perform a full twist (https://youtu.be/tfY6_C5SswU). In our sensitivity analysis, we found at least two origins of the ‘twist-only’ behaviour across smaller sample sizes (N=2-10), but there were not enough species with large numbers of observations (N>10) to reconstruct the evolution of the trait (Supplementary Fig. 1). The two stable origins were consistently found in *Dicranosepsis unipilosa* and at the *Sepsis* clade, but higher numbers of acquisition (4-6) were found when all species were analysed (four for N=6-7, six for N=8-10; Supplementary Figure 1).

**Figure 2.**
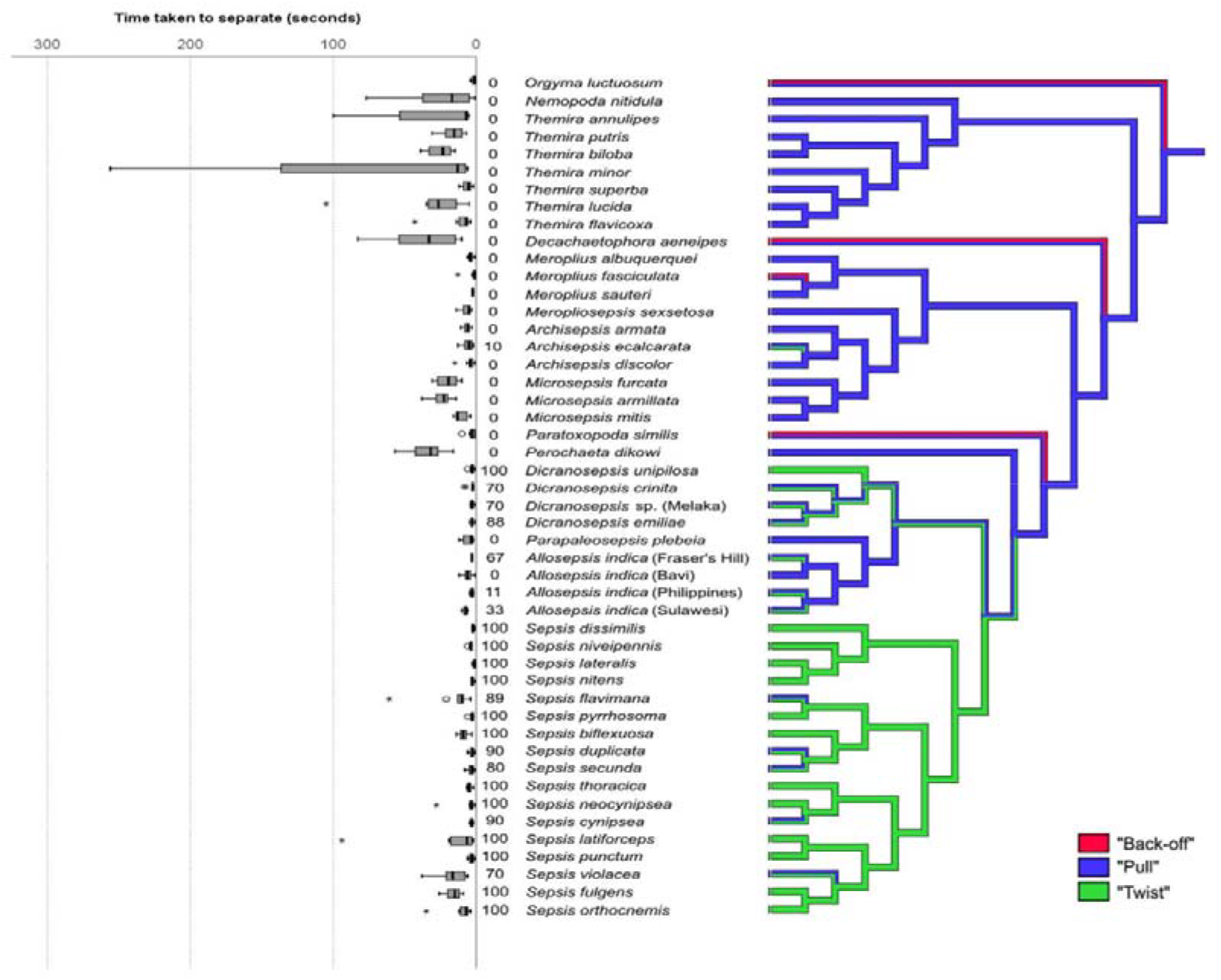
Evolution of separation behaviours and times. Right panel: Ancestral state reconstructions of the three types of separation behaviour (‘back off’, ‘pull’, ‘twist’). Percentage of trials with ‘twists’ indicated at base of boxplot. Left panel: Separation time.

### Relationship between separation technique and sexual size dimorphism

The mean SSD ratios for ‘pull’ males were only slightly higher (‘pull’: 0.95±0.11, N=24; ‘twist’: 0.90±0.09, N=36). We did not find any significant effect of size in explaining whether larger males in species with intraspecific variation in separation were inclined to perform a ‘pull’ or a ‘twist’ (logistic regression; X^2^=3.15, d.f.=1, p=0.076).

### Relationship between separation technique, time and variance

Separation times of males in those species that do not ‘twist’ were significantly longer and the variance was significantly higher (‘pull’ only: 545.37±1953.94 s, N=164; ‘twist’/ ‘pull’: 164.29±228.75, N=213; Brown-Forsythe test: F_(1, 362)_ = 5.4118, p = 0.02; Welch test: F_(1, 161.33)_ = 5.55, p=0.02) suggesting that twisting produces shorter and less variable separation times. When accounting for phylogenetic relatedness, neither SSD nor separation behaviour were significant predictors for separation time (final model: separation time = 3.6209 - 4.5954 * SSD ratio, N=33, R^2^=0.04, F_(1, 31)_=1.314, p=0.26). However, it is important to remember the limitations to using the phylogenetic generalized linear model due to its reliance on averaged data with high standard deviations instead of utilizing all individual data points.

### Handedness

There were clear, observable species-specific differences in phallus morphology and symmetry across sepsids (Fig. 3–5). Although, there was no significant relationship between technique employed and directionality of separation (X^2^=5.63, d.f.=2, p=0.06), but we observed a positive trend (81-90%) within *Microsepsis* (3 species, N=10-14) where males were pulling from the females in a clockwise direction (Fig. 5).

**Figure 3.**
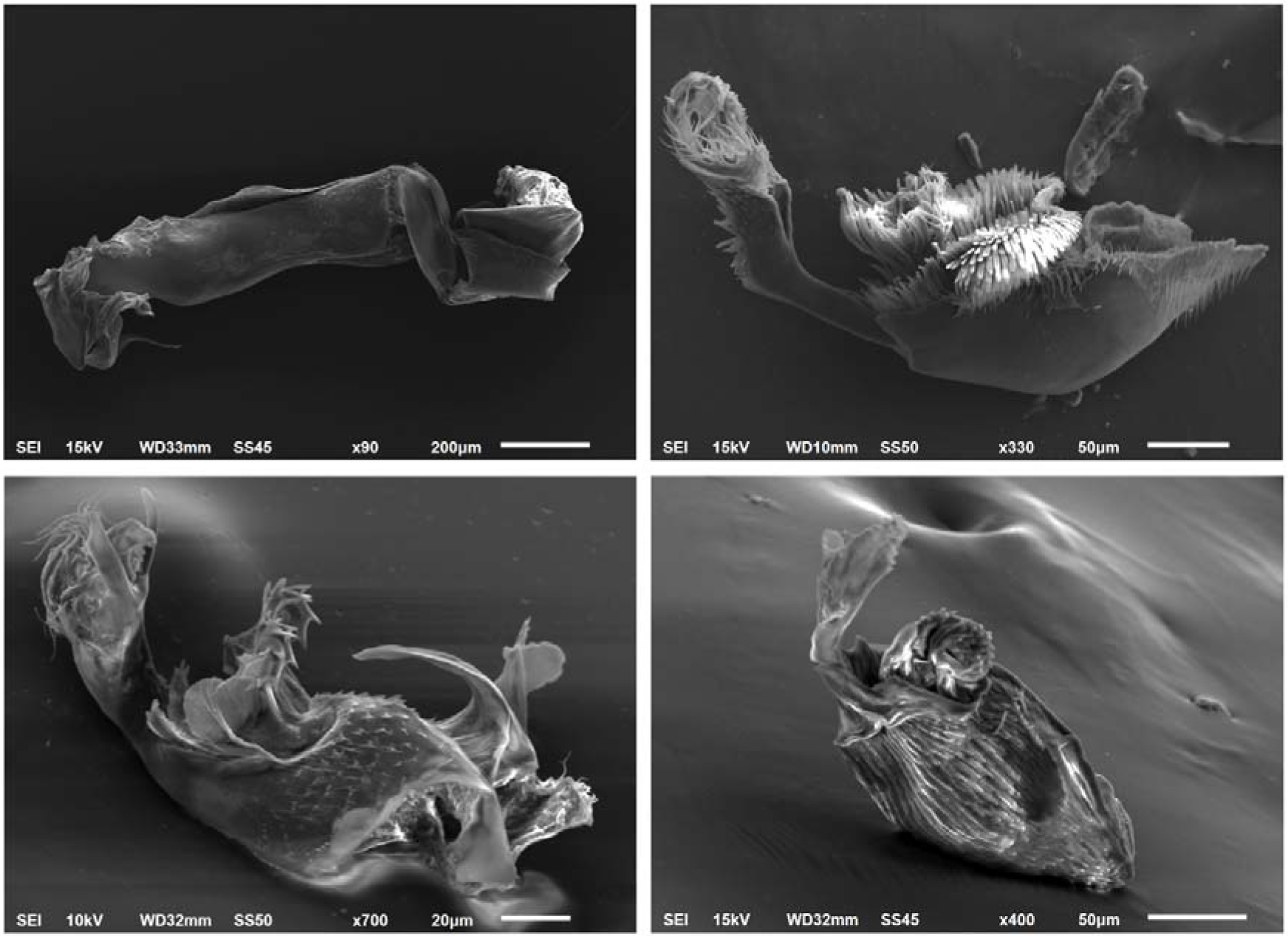
Phallus morphology of sepsid species with different separation techniques; A) *Orgyma luctuosum* (‘back-off’, ‘pull’), B) *Nemopoda nitidula* (‘pull’), C) *Sepsis dissimilis* (‘twist’), D) *Dicranosepsis crinita* (‘pull’ and ‘twist’).

## Discussion

The termination of copulation is an event that can have fitness consequences for both sexes. In some species, this event can be fraught with conflict, where males attempt to maintain genital contact beyond the copulation time preferred by the female (e.g. *Drosophila montana*; Mazzi et al. 2009). In others, the termination of copulation ends with a nuptial gift (e.g. Vahed 1998). However, it is rare to find that studies that characterize the separation behaviour in detail, let alone document inter- and intraspecific variation with regard to separation behaviour. This is the first comparative analysis of separation behaviour across a sizeable insect clade. We find that male sepsids employ three different techniques to break genital contact with females. The most conspicuous technique is ‘twist’, where the males use the genital contact with females as a ‘hinge’ to ‘twist’ around in either clock- or anti-clockwise direction to facilitate separation. We show both quantitatively and qualitatively, a clear shift from ‘pull’ as the ancestral mode of separation in Sepsidae to ‘twist’ as a derived trait. There are apparently multiple origins of ‘twist’ in Sepsidae (Fig. 2), and a long phase of transitioning to ‘twist’ with many species being polymorphic in that they have both ‘pull’ and ‘twist’ in their repertoire (12 species). ‘Twist’ is likely fixed in 13 species (N≥4).

Our qualitative results suggest that intraspecific variation in separation behaviour may be a result of sepsid males using the default technique first and then trying other movements when the default leads to protected separation times. This is observed in “pull” species where males that struggle to separate seem to start improvising by wiggling their abdomen and moving to the left or right instead of just heading straight in the opposite direction (*N. nitidula*: https://youtu.be/tfY6_C5SswU). In another species, the male chances upon the ‘twist’ after he failed to separate from the female via ‘pull’ (*A. ecalcarata*: https://youtu.be/uNPB7cURU7A). Although ‘twist’ was a fixed trait for 13 species, the number of revolutions a male performs in a ‘twist’ is not fixed within a species. This suggests that most males in the *Sepsis* group (Meier, 1995; 1996; Su et al., 2008; Lei et al., 2013) have the flexibility of using either ‘pull’ or ‘twist’ in order to separate in a fast and predictable manner. Males may use ‘pull’ first but then change to ‘twist’ when separation is not occurring quickly enough. This may explain the observation of intraspecific variation in some species.

‘Twist’ only species may have evolved such behaviour because it increases termination efficiency: shorter, and less variable separation time (Fig. 2). The reduced variation in the time taken to separate may be the adaptive significance of the ‘twist’ that has led to its eventual fixation in some species. Short separation times can affect fitness by reducing exposure to predation or reducing the refractory period between copulations (Wing 1988; Rowe 1994; Hosken et al. 1994; Kotiaho et al. 1998; Kemp 2012; Siemers et al. 2012; but see McCauley and Lawson 1986). This also applies to mating pairs that are struggling to break genital contact. Furthermore, sepsids that are in-copula can perform short bursts of flight, led by the female, but this is not an option for pairs that are struggling to separate (pers. obs). As such, this vulnerable end-phase of copulation causes an even greater predation risk than copulation per se.

One important factor to consider would be the relationship between genitalia morphology and separation technique. This requires morphological data for the male phallus (Fig. 3) and the female reproductive tract. Unfortunately, both contain soft and membranous parts and the sepsid female internal reproductive tract, in particular, has little sclerotization (Eberhard and Huber 1998; Puniamoorthy et al. 2010). This makes it difficult to study co-evolution with reasonable precision. In addition, a detailed description of male-female genitalic interactions in *Archisepsis* (Eberhard and Huber, 1998) finds intraspecific variability in contact interactions, which could be due to the flexibility of the female tract. Additionally, the membranous cover of non-interlocking sclerotized plates of the male phallus are also difficult to quantify (Araujo 2015). A recent study of sepsid phallus morphology grouped species according to phallus similarity but these groups are not correlated with separation technique (Araujo 2015; see https://sketchfab.com/daedalus for 3D models of aedeagi from these clades). Asymmetrical phallus are common in sepsid species (Araujo 2015) but the directionality of separations are random for almost of the species studied in this study. The only exception is a positive trend in the three species in *Microsepsis* (Suppl.; Figure C). Note that the surstyli in these species are particularly asymmetrical (external genitalic clasping organs) and that surstyli play a role in gaining access to female genitalic tract opening and intromission.

Copulatory wounding can occur when the intromittent organs of males are armoured and have spiny protrusions (Merritt 1989; Crudgington and Siva-Jothy 2000; Allard 2006; Kamimura 2007, Okuzaki et al. 2012). The only detailed studies of such interactions in sepsids are of three *Archisepsis* species, which tend to have forceful interactions between male and female structures throughout copulation. In fact in *A. armata*, the long finger of the male intromittent organ likely results in the post-ejaculation deformation of the female vagina (Eberhard and Huber 1998). If these injuries were a consequence of extraneous tugging during separation, then specific areas that are in contact during separation that would most likely have damage. However most *Archisepsis* do not have difficult separations and generally have short separation times (Fig. 1). A possible avenue for future research would be to investigate detailed interaction in species with slow separation (e.g., *Nemopoda nitidula*, *Decachaetophora aeneipes*, *Perochaeta dikowi*) as well as species where complete separation failure has been observed regularly (*Sepsis cynipsea*, *Themira biloba*).

Finally, we note that the complex genitalia of most sepsids differ considerably from the aedeagus of *Orygma luctuosum*, which may be the sister species of all remaining sepsids and possesses a smooth, non-spiny aedeagus. *Orygma* is a fast separator which uses ‘back-off’ and ‘pull’ (Fig. 3–4). Despite having spiny armoured aedeagi, species in *Nemopoda*, *Themira* and *Decachaetophora* continue to use ‘pull’ or ‘back-off’, but many species in these genera have long separation times (Fig. 1). It is conceivable that ‘twist’ could have emerged as a solution to separation difficulties that evolved because of spiny aedeagi. Such a scenario would be consistent with the observed shift to ‘twist’ and a reduction in separation time in the *Sepsis* group. It is also plausible that the genital rotation occurring during ‘twist’ may aid in dislodgement of the genital spines, “anti-slip” devices and “anchors” (Arauja 2015).

**Figure 4.**
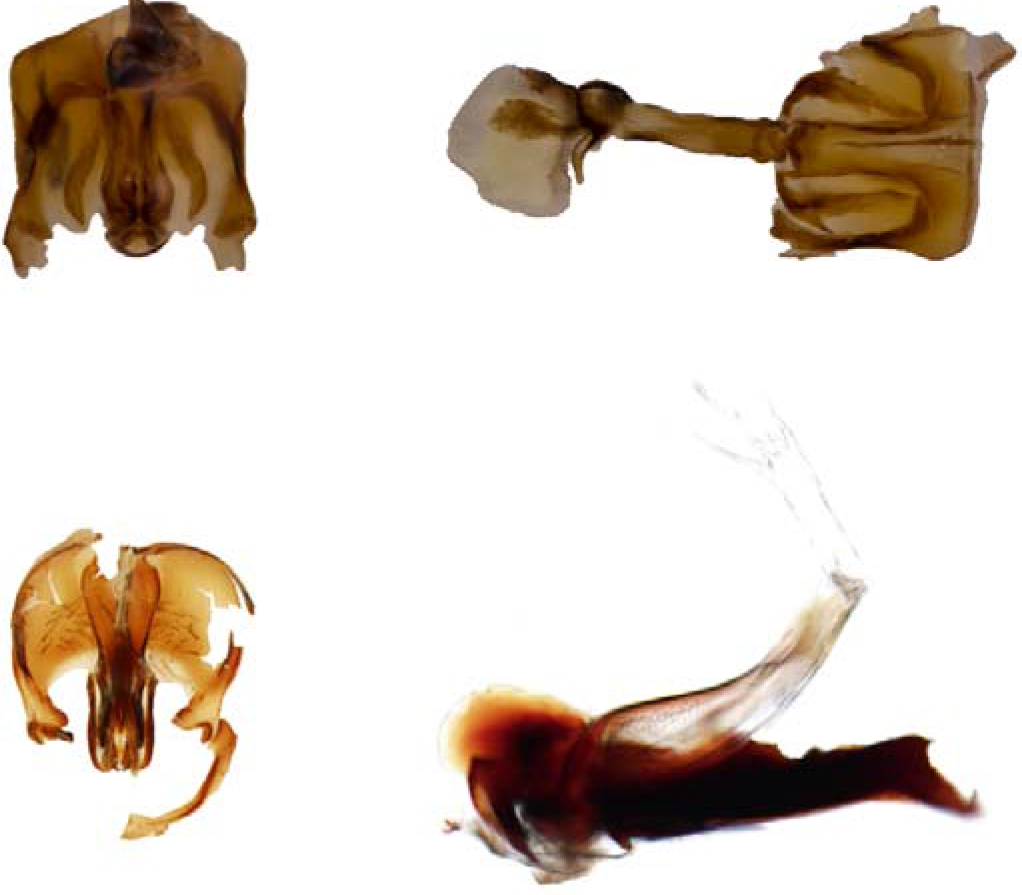
Phallus morphology of *Orygma luctuosum* (top) and *Ortalischema albitarse* (bottom). A, C) Dorsal view of hypandrium with phallus resting inside of it. B) Ventral and D) lateral view of phallus.

**Figure 5.**
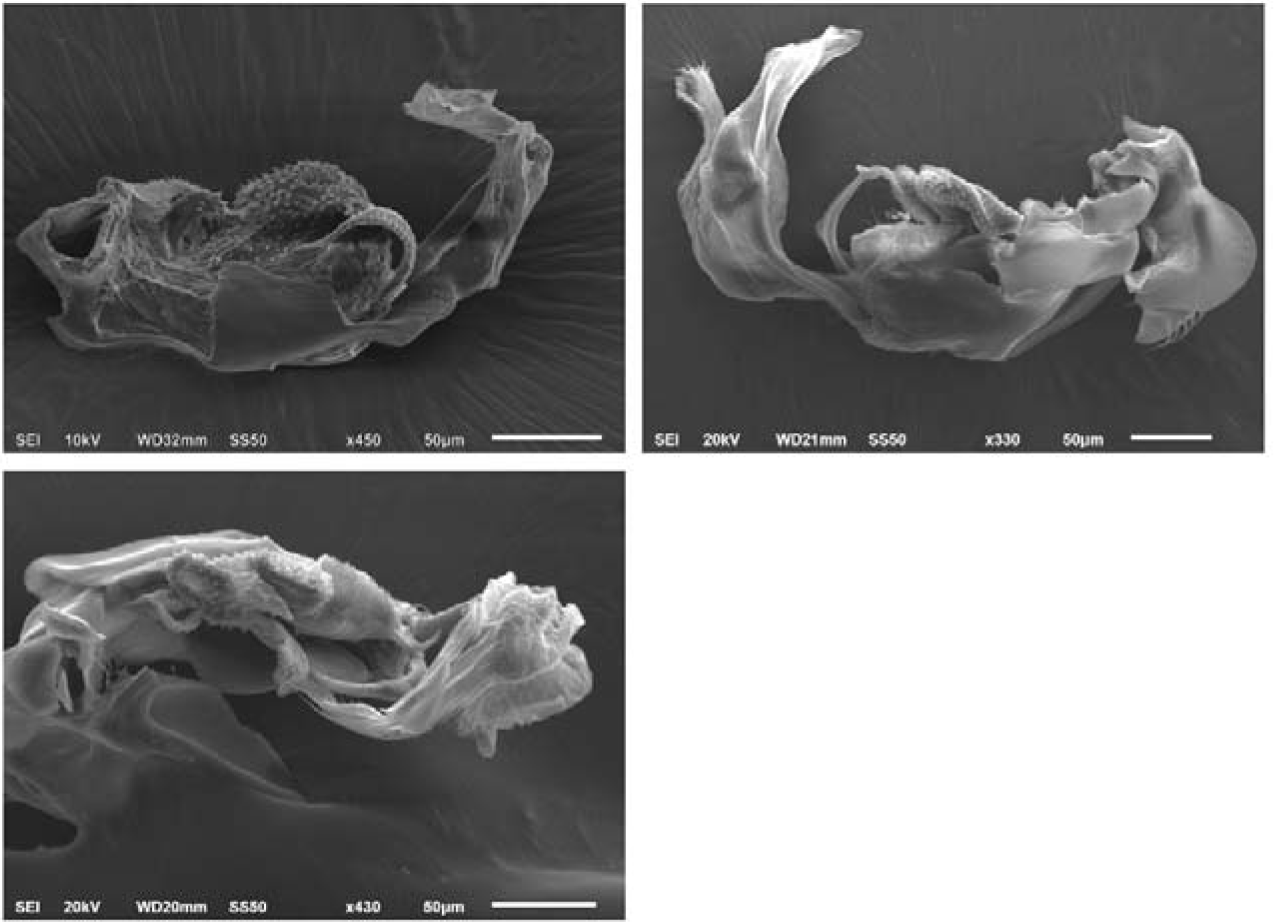
Phallus morphology of 3 species of *Microsepsis*. A) *Microsepsis furcata*, B) *Microsepsis armillata*, C) *Microsepsis mitis*.

In summary, our study reveals a surprising diversity of separation behaviours both within and across 48 species. The conspicuous ‘twist’ evolved multiple times, possibly in order to increase the efficiency of separation, for example, for reducing the risk of predation. We found no direct correlation between male genital morphology and separation behaviour, but there are numerous practical challenges to studying such interactions because they involve membranous parts of male and female genitalia. Future studies of separation behaviour could focus on species that exhibit intraspecific variation in separation techniques to assess whether such differences in ‘pull’ versus ‘twist’ result in differential fitness (e.g. male/female survival, re-mating, offspring etc.).

## Acknowledgements

We thank numerous students from the Evolutionary Biology Laboratory at the National University of Singapore for recording and evaluating the behaviour.. RM would like to acknowledge support from MOE Grant R-154-000-A62-112.

## References

Allard D., Børgesen L., Van Hulle M., Bobbaers A., Billen J. and Gobin B. 2006. Sperm transfer during mating in the pharaohs ant, *Monomorium pharaonis*. Physiol. Entomol. 31: 294–298.

Ang Y.C., Puniamoorthy, J., Pont A. C., Bartak, M., Blanckenhorn, W. U., Eberhard, W. G., Puniamoorthy, N., Silva, V., Munari, L., Meier, R. 2013. SepsidNet: A plea for digital reference collections and other science-based digitization initiatives in taxonomy: Sepsidnet as exemplar. Systematic Entomology 38: 637–644.

Araujo, D.P. 2015. The virtual biology of sepsid flies: 3D computer graphics tools for research in morphology, systematics and biomechanics. PhD dissertation. National University of Singapore.

Arnqvist G. and I. Danielsson. 1999. Copulatory behavior, genital morphology and male fertilization success in water striders. Evolution 53:147–156.

Baer B. and J.J. Boomsma. 2006. Mating biology of the leaf-cutting ants *Atta colombica* and *A. cepha*lotes. J. Morph. 267: 1165–1171.

Blanckenhorn W.U., Hosken D.J., Martin O.Y., Reim C., Teuschl Y. and P.I. Ward. 2002. The costs of copulating in the dung fly *Sepsis cynipsea*. Behavioural Ecology 13(3):353–358.

Boake C.R.B. 1989. Repeatability: its role in evolutionary studies of mating behavior. Evolutionary Ecology 3:173–182.

Bonduriansky R. 2003. Layered sexual selection: a comparative analysis of sexual behaviour within an assemblage of piophilid flies. Can. J Zool 81:479–491.

Brown, M. B., and A. B. Forsythe. 1974. The small sample behavior of some statistics which test the equality of several means. Technometrics, 16, 129–132.

Cook D.F. Differences in courtship, mating and postcopulatory behaviour between male morphs of the dung beetle *Onthophagus binodis* Thunberg (Coleoptera: Scarabaeidae). Animal Behaviour 40:428–436.

Crudgington H.S. and M.T. Siva-Jothy. 2000. Genital damage, kicking and early death. Nature 407:855–856.

Eberhard, W.G. 1985. Sexual Selection and Animal Genitalia. Harvard University Press.

Eberhard W.G. 1998. Reproductive behavior of *Glyphidops flavifrons* and *Nerius plurivitatus* (Diptera, Neriidae). Journal of the Kansas Entomological Society 71(2):89–107.

Eberhard W.G. 2001. Species-specific genitalic copulatory courtship in sepsid flies (Diptera, Sepsidae, *Microsepsis*) and theories of genitalic evolution. Evolution 55(1):93–102.

Eberhard W.G. 2002. The relation between aggressive and sexual behavior and allometry in *Palaeosepsis dentatiformis* flies (Diptera: Sepsidae). Journal of the Kansas Entomological Society 75(4):317–332.

Eberhard W.G. 2008. Static allometry and animal genitalia. Evolution 63(1):48–66.

Eberhard, W.G. and F. Pereira. 1996. The functional morphology of male genitalic surstyli in the dungflies *Achisepsis diversiformis* and *A. ecalcarata*. J. Kans. Entomol. Soc. 69, suppl.: 43–60.

Eberhard W.G. and B.A. Huber. 1998. Copulation and sperm transfer in *Archisepsis* flies (Diptera, Sepsidae) and the evolution of their intromittent genitalia. Studia dipterologica 5(2):217–248.

Eberhard W.G., Bernhard B., Huber B.A., Rodriguez R., Briceño D. Salas I. and V. Rodriguez. 1998. One size fits all? Relationships between the size and degree of variation in genitalia and other body parts in twenty species of insects and spiders. Evolution 52:415–431.

Fox J and S. Weisberg. 2011. An {R} Companion to Applied Regression, Second Edition. Thousand Oaks CA: Sage. URL: http://socserv.socsci.mcmaster.ca/jfox/Books/Companion

Grafen A. 1989. The phylogenetic regression. Philos. Trans. R. Soc. London Ser. B 326:119–57.

Herberstein M.E., Wignall A.E., Nessler S.H., Harmer A.M.T. and Schneider J.M. 2012. How effective and persistant are fragments of male genitalia as mating plugs? Behav. Ecol. 1140–1145.

Hosken D.J., Bailey W.J., O’Shea J.E. and J.D. Roberts. 1994. Localization of insect calls by the bat *Nyctophilus geoffroyi* (Chiroptera: Vespertilionidae): a laboratory study. Aust J Zool 42 : 177–184.

Kamimura Y. 2007. Twin intromittent organs of *Drosophila* for traumatic insemination. Biol. Lett. (2007) 3, 401–404.

Kamimura Y. 2008. Copulatory wounds in the monandrous ant species *Formica japonica* (Hymenoptera: Formicidae). Ins Soc 55:51–53.

Kemp D.J. 2012. Costly copulation in the wild: mating increases the risk of parasitoidmediated death in swarming locusts. Behav. Ecol. 23, 191–194.

Kotiaho J.S., Alatalo R.V., Mappes J., Parri S. and A. Rivero. 1998. Male mating success and risk of predation in a wolf spider: a balance between sexual and natural selection? J. Anim. Ecol. 67: 287–291.

Kummer H. 1960. Experimentelle Untersuchungen zur Wirkung von Fortpflanzungs-Faktoren auf die Lebensdauer von *Drosophila melanogaster* Weibchen. Z Vgl Physiol 43:642–679.

Lei Z., Ang S.H.A., Srivathsan A., Su F-Y.K. and R. Meier 2013. Does better taxon sampling help? A new phylogenetic hypothesis for Sepsidae (Diptera: Cyclorrhapha) based on 50 new taxa and the same old mitochondrial and nuclear markers. Molecular Phylogenetics and Evolution 69:153–164.

Maddison W.P. 1990. A method for testing the correlated evolution of two binary characters: are gains or losses concentrated on certain branches of a phylogenetic tree? Evolution: 44(3):539–557.

Maddison W.P. 2000. Testing character correlation using pairwise comparisons on a phylogeny. J. Theoretical Biology. 202: 195–204.

Maddison W.P. and R.G. Fitzjohn. 2015. The unsolved challenge to phylogenetic correlation tests for categorical characters. Systematic Biology 64:127–136.

Maddison W. P. and D.R. Maddison. 2015. Mesquite: a modular system for evolutionary analysis. Version 3.04 http://mesquiteproject.org

Martins E.P. and T.F. Hansen. 1997. Phylogenies and the comparative method: a general approach to incorporating phylogenetic information into the analysis of interspecific data. Am. Nat. 149:646–67.

Mayr E. 1963. Animal Species and Evolution. Harvard University Press, Cambridge, Massachusetts.

Mazzi D., Kesäniemi J., Hoikkala A. and K. Klappert. 2009. Sexual conflict over the duration of copulation in Drosophila montana: why is longer better? BMC Evol. Biol. 9: 132.

McCauley D.E. and E.C. Lawson. 1986. Mating reduces predation on male milkweed beetles. Am. Nat. 127: 112–117.

Meier, R. 1995. Cladistic analysis of the Sepsidae (Cyclorrhapha: Diptera) based on a comparative scanning electron microscopic study of larvae. Systematic Entomology 20: 99–128.

Meier, R. 1996. Larval morphology of the Sepsidae (Diptera: Sciomyzoidea), with a cladistic analysis using adult and larval characters. Bulletin of the American Museum of Natural History 228: 1–147.

Meier, R. 2017. Citation of taxonomic publications: the why, when, what, and what not. Systematic Entomology 42: 301–304.

McCullagh P. and J.A. Nelder. 1997. Generalized Linear Models: Monographs on Statistics and Applied Probability. Chapman and Hall, London.

Merritt D.J. 1989. The morphology of the phallosome and accessory gland material transfer during copulation in the blowfly, *Lucilia cuprina* (Insecta, Diptera). Zoomorphology 108:359–366.

Okuzaki Y., Takami Y., Tsuchiya Y. and T. Sota. 2012. Mating behavior and the function of the male genital spine in the ground beetle *Carabus clathratus*. Zoological Science 29(7):428–432.

Orme D., Freckleton R., Thomas G., Petzoldt T., Fritz S., Isaac N. and W. Pearse. 2013. caper: Comparative analyses of phylogenetics and evolution in R. R package version 0.5.2. http://CRAN.R-project.org/package=caper

Pagel M. 1994. Detecting correlated evolution on phylogenies: a general method for the comparative analysis of discrete characters. Proc. R. Soc. London B 255: 37–45.

Paradis E., Claude, J. and Strimmer, K. 2004. APE: analyses of phylogenetics and evolution in R language. Bioinformatics 20: 289–290.

Parker G.A. 1972. Reproductive behaviour of *Sepsis cynipsea* (L.) (Diptera: Sepsidae) I. A preliminary analysis of the reproductive strategy and its associated behaviour patterns. Behaviour 41: 172–206.

Pont, A. C., and R. Meier. 2002. The Sepsidae (Diptera) of Europe. Fauna Entomologica Scandinavica 37: 1–221.

Puniamoorthy, N., K. Feng Yi Su, R. Meier. 2008. Bending for love: losses and gains of sexual dimorphisms are strictly correlated with changes in the mounting position of sepsid flies (Sepsidae: Diptera). BMC Evolutionary Biology 8:155.

Puniamoorthy N., Ismail M.R., Tan D.S. and R. Meier. 2009. From kissing to belly stridulation: comparative analysis reveals surprising diversity, rapid evolution, and much homoplasy in the mating behaviour of 27 species of sepsid flies (Diptera: Sepsidae). Journal of Evolutionary Biology 22(11):2146–2156.

Puniamoorthy N., Kotrba M. and R. Meier. 2010. Unlocking the “black box“: internal female genitalia in Sepsidae (Diptera) evolve fast and are species-specific. BMC Evolutionary Biology 10:275.

R Core Team. 2013. R: A language and environment for statistical computing. R Foundation for Statistical Computing, Vienna, Austria. ISBN 3-900051-07-0, URL http://www.R-project.org/.

Read A.F. and S. Nee. 1995. Inference from binary comparative data. J. Theoretical Biology 173:99–108.

Ross K.G. 1983. Laboratory studies of the mating biology of the eastern yellowjacket, *Vespula maculifrons* (Hymenoptera: Vespidae). Journal of the Kansas Entomological Society 56(4):523–537.

Rowe L. 1994. The costs of mating and mate choice in water striders. Anim. Behav. 48, 1049–1056.

Siemers B.M., Kriner E., Kaipf I., Simon M. and S. Greif. 2012. Bats eavesdrop on the sound of copulating flies. Curr. Biol. 22:R563–R564.

Spieth H.T. 1974. Courtship behavior in *Drosophila*. Annual Rev. Entomol. 19:385–405.

Su Feng Yi, K., S. Kutty and R. Meier 2008. Morphology versus Molecules: The phylogenetic relationships of Sepsidae (Diptera: Cyclorrhapha) based on morphology and DNA sequence data from ten genes. Cladistics 24: 902–916.

Tan D.S.H., Ng S.R. and R. Meier. 2011. New information on the evolution of mating behaviour in Sepsidae (Diptera) and the cost of male copulations in *Saltella sphondylii*. Organisms Diversity & Evolution 11(4):253–261.

Vahed, K. (1998). The function of nuptial feeding in insects: a review of empirical studies. Biol. Rev. 73, 43–78.

von Helversen D. and O. von Helversen. 1991. Pre-mating sperm removal in the bushcricket *Metaplastes ornatus* Ramme 1931 (Orthoptera, Tettigonoidea, Phaneropteridae). Behav. Ecol. Sociobiol. 28: 391–396.

Wang Q., Chen L., Li J. and X Yin. 1996. Mating behavior of *Phytoecia rufiventris* Gautier (Coloeptera: Cerambycidae). Journal of Insect Behavior 9(1):47–60.

Welch B.L.. 1951. On the comparison of several mean values: an alternative approach. Biometrika 38, 330–336.

West-Eberhard M.J. 1984. Sexual selection, competitive communication and species-specific signals in Insects. In: Insect Communication (ed. T. Lewis) pp. 283–324. London: Academic Press.

Wojcik D.P. 1969. Mating behavior of 8 stored-product beetles (Coleoptera: Dermestidae, Tenebrionidae, Cucujidae, and Curculionidae). The Florida Entomologist 52(3):171–197.

Wing S.R. 1988. Cost of mating for female insects: risk of predation in *Photinus collustrans*. Am. Nat. 131, 139–142.

